# Compact Standalone Platform for Neural Recording with Real-Time Spike Sorting and Data Logging

**DOI:** 10.1101/186627

**Authors:** Song Luan, Ian Williams, Michal Maslik, Yan Liu, Felipe de Carvalho, Andrew Jackson, Rodrigo Quian Quiroga, Timothy G. Constandinou

## Abstract

**Objective:** Longitudinal observation of single unit neural activity from large numbers of cortical neurons in awake and mobile animals is often a vital step in studying neural network behaviour and towards the prospect of building effective Brain Machine Interfaces (BMIs). These recordings generate enormous amounts of data for transmission & storage, and typically require offline processing to tease out the behaviour of individual neurons. Our aim was to create a compact system capable of: 1) reducing the data bandwidth by circa 3 orders of magnitude (greatly improving battery lifetime and enabling low power wireless transmission); 2) producing real-time, low-latency, spike sorted data; and 3) long term untethered operation. *Approach*. We have developed a headstage that operates in two phases. In the short training phase a computer is attached and classic spike sorting is performed to generate templates. In the second phase the system is untethered and performs template matching to create an event driven spike output that is logged to a micro-SD card. To enable validation the system is capable of logging the high bandwidth raw neural signal data as well as the spike sorted data. *Main results*. The system can successfully record 32 channels of raw neural signal data and/or spike sorted events for well over 24 hours at a time and is robust to power dropouts during battery changes as well as SD card replacement. A 24-hour initial recording in a non-human primate M1 showed consistent spike shapes with the expected changes in neural activity during awake behaviour and sleep cycles. *Significance* The presented platform allows neural activity to be unobtrusively monitored and processed in real-time in freely behaving untethered animals revealing insights that are not attainable through scheduled recording sessions and provides a robust, low-latency, low-bandwidth output suitable for BMIs, closed loop neuromodulation, wireless transmission and long term data logging.

## 1. Introduction

There is currently a global effort to expand our knowledge of the brain by recording large-scale cortical neural activity over a long period of time. The ‘Brain Research through Advancing Innovative Neurotechnologies’ (BRAIN) Initiative in particular is looking to deliver a step change in our understanding of the brain’s networks and function – a key element of this is research to advance neural recording from the current state-ofthe-art (short term targeted recordings of in the order of 1000s of channels [1, 2, 3]) to round-the-clock recordings of more than 100,000 channels in freely behaving animals [4].

These ever increasing channel counts and recording periods naturally lead to an enormous amount of raw data to be stored and processed to extract useful information. For example a 100,000 channel system could well generate in excess of 10 Gigabits per second of data, not to mention the requirement of recording 24 hours. This presents a major obstacle for chronic operation and real-time behaviour analysis. However, raw recorded neural data is sparse and hence amenable to data compression [5, 6, 7] or techniques to extract useful information from the signal locally on the recording device [8, 9, 10]. While both these methods are effective in saving bandwidth and power [11], the latter also enables on-node network state analysis and triggers for controlling brain machine interfaces (BMI) without data decompression.

Understanding network state and correlating it with observed or desired behaviour is greatly enhanced by separating out the activity of individual neurons from the melee recorded by each electrode – i.e. spike sorting [12, 13] – as each neuron may encode different features of a behaviour. Naive spike sorting (involving spike detection and clustering) is a computationally complex process that is normally performed offline on a computer using Principal Component Analysis or wavelet decomposition. However, in recent years a two step process has emerged: in the first step naive sorting is performed offline to identify characteristics of the spikes; and in the second step a computationally simpler form of spike sorting can be performed online based on classification rather than clustering. These classification methods include: feature extraction based on first derivative peaks [8]; or zero crossings [14]; or key differentiating points [15]. The system presented here utilises template matching to achieve high accuracy with exceptionally low computational cost [1, 16, 17] making it attractive for high channel count systems.

This paper describes a system (see Figure 1) capable of recording raw neural signal data and performing real-time template matching based spike sorting. The event driven spike sorted output reduces the data transmission bandwidth by circa 3 orders of magnitude (assuming a nominal spike rate of 10 spikes per second [18]) and provides a sub-millisecond latency control signal for closed-loop neuromodulation or brain machine interfaces.

**Figure 1.**
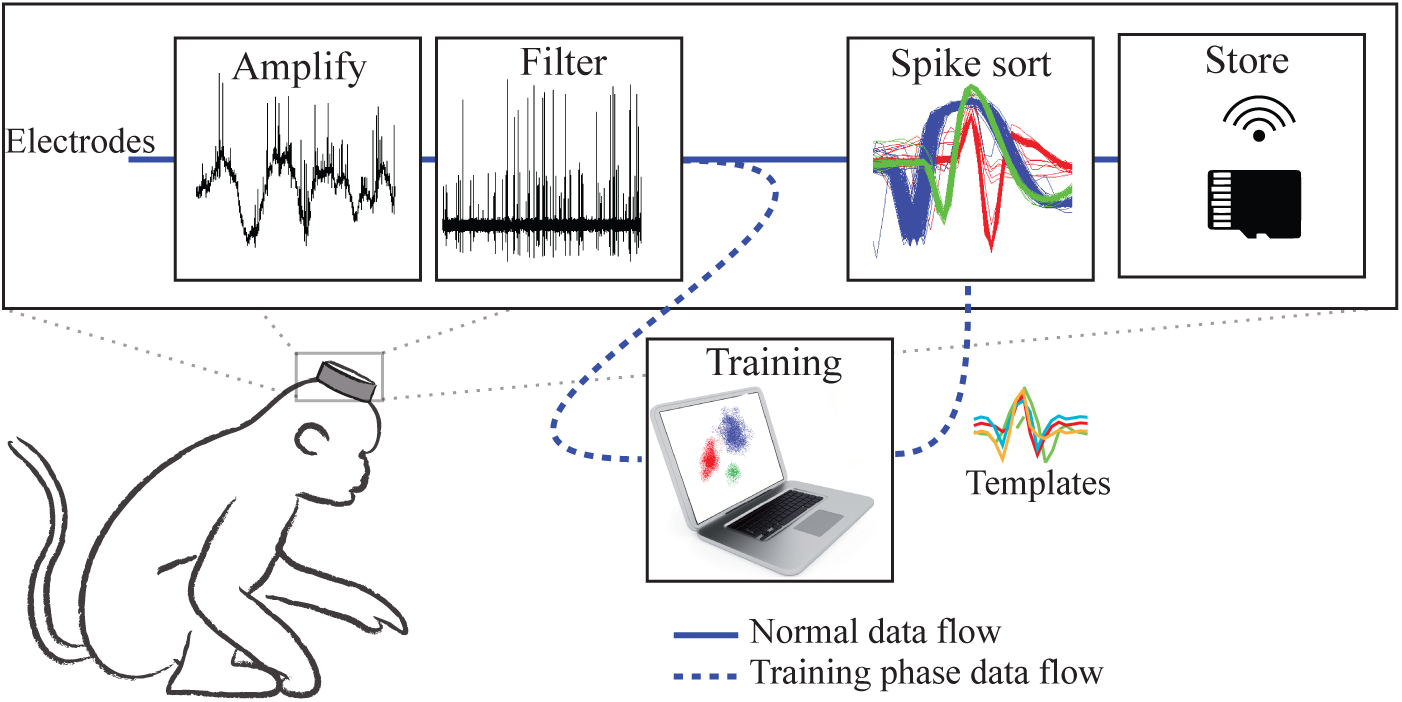
System concept – the miniature head-mounted system amplifies, filters and spike sorts neural signals from electrodes inserted in the cortex, and the resulting spike events are either stored in local memory (as demonstrated here) or wirelessly transmitted. During a training phase a cable to a computer is briefly attached and raw neural signal data is recorded, clustered and the resulting characteristic spike waveforms are uploaded into the headstage to enable the template matching based spike sorting.

A similar concept was demonstrated by Schwarz *et al.* [1], however, they identified a bidirectional wireless link as essential for on-node spike sorting and as a result their system was limited to only logging 6 channels of neural signal (plus spike sorted events) due to the limited wireless bandwidth available and also suffered from wireless transmission latency and packet loss. Here we present a system capable of lossless recording of 32 channels of neural signal by using a two phase approach to spike sorting as well as on-node data logging. Further advantages of this system include: the modular rather than monolithic design (which enables the on-node data logging to be easily replaced by a wireless transceiver or BMI); a more compact front end and spike sorting module; an increase in the number of templates per channel (and hence distinguishable neurons) from 2 to 4; and features targeted at chronic experiments such as automatic reloading of configuration data on power cycling or SD card changing. The system has been fully tested *in-vivo* for several 24-hour recording sessions.

The remainder of this paper is organised as follows: Section 2 describes the system design and implementation; Section 3 presents performance metrics from benchtop electronic testing as well as the results obtained from *in-vivo* experiments; and Section 4 discusses the results, limitations and expected future development of the system.

## 2. System Design

### 2.1. Outline system architecture

The system (shown in Figure 2) has been designed in a modular fashion with standard connectors to enable future flexibility and system reconfiguration. The core of the system is the Neural Interface board which performs the signal conditioning and on-node spike sorting. For initial system validation this has been paired with a custom micro-SD logging and standalone controller board capable of recording all the raw and spike sorted data as well as automatic reconfiguring the system when power is interrupted (e.g. during a battery change) or when the SD card is changed. These two boards have been designed to be small and lightweight to enable them to be head-mounted for long periods of time. In addition a benchtop Data Interface board can be connected through a wired link to enable data to be logged to a computer and for configuration data to be uploaded to the head-mounted boards.

**Figure 2.**
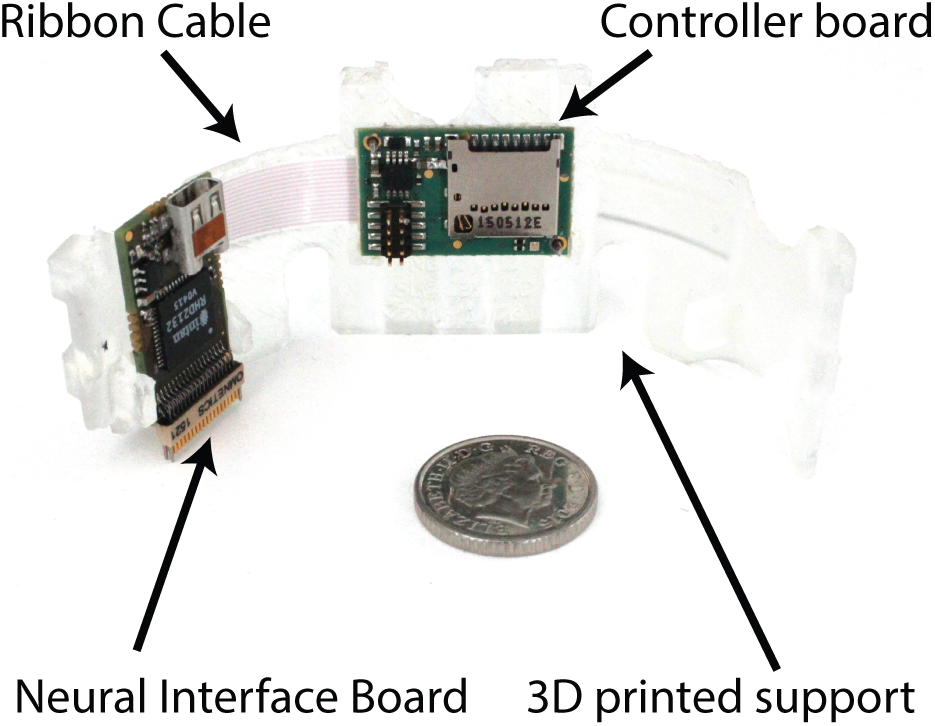
The standalone system mounted in a 3D printed support designed to fit inside a monkey’s head cradle.

### 2.2. Typical system operation

The system is designed to operate in two distinct phases of operation as shown in Figure 3.

**Figure 3.**
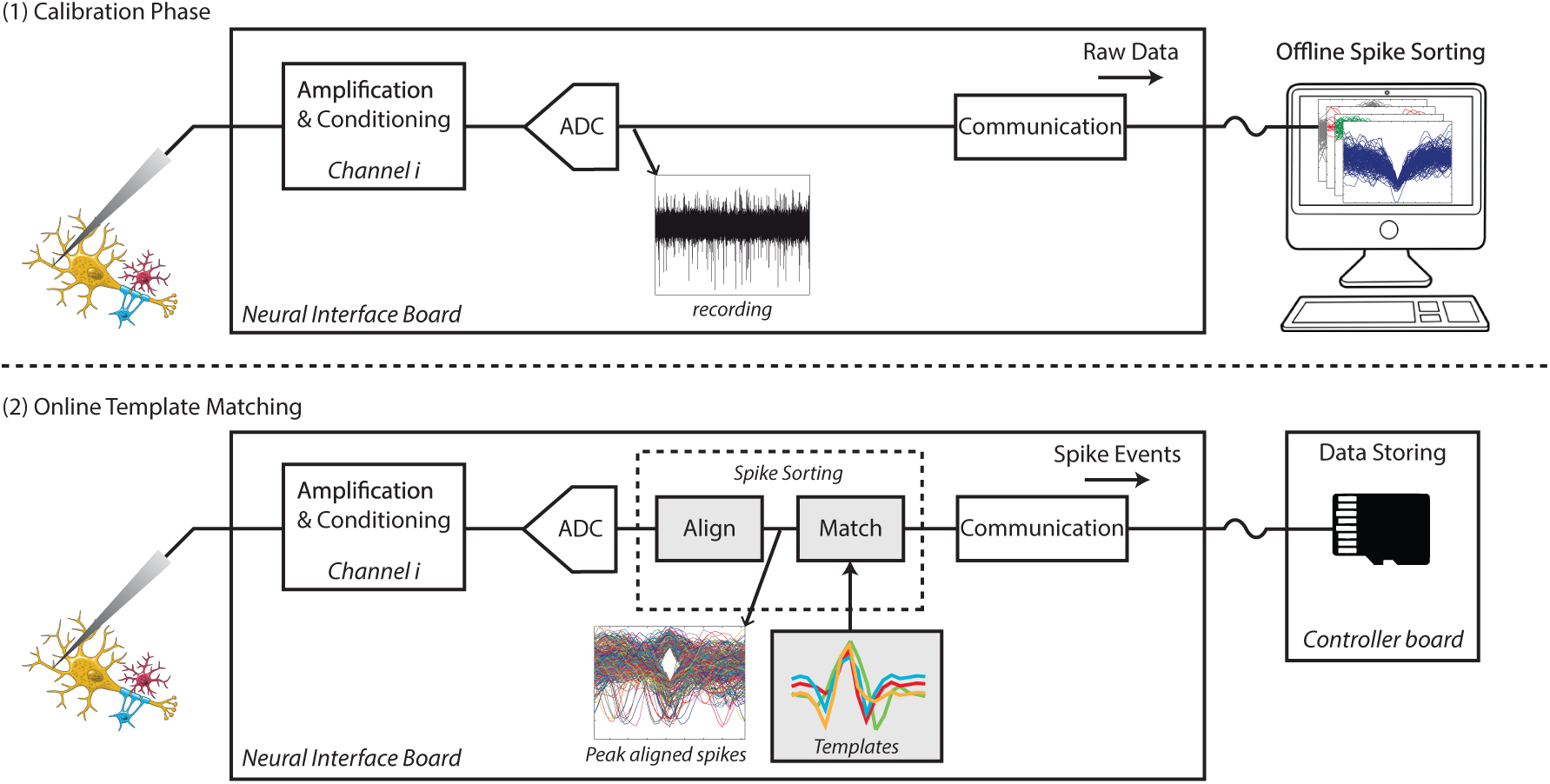
Strategy of two-phase online spike sorting. (1) Calibration phase using a computer to generate templates offline. (2) Real-time on-node template matching where spike events are stored on an SD card.

i. In the calibration phase, the head-mounted system is tethered to a computer and a 5-10 minute recording of raw neural signal data is recorded and clustered using offline processing (here we use Wave Clus [19]). Spike detection levels, characteristic templates for each of the observed spike clusters, and template matching thresholds are then uploaded for the next phase.
ii. During the online template matching phase, the headstage is untethered from the computer and a small, low-power Field Programmable Gate Array (FPGA) is used to perform template matching based spike sorting. The resulting raw and/or spike sorted data is recorded to a micro-SD card for later analysis and validation.

### 2.3. System design rationale

Real-time on-node template matching lies at the heart of this system and typically involves 3 main processing steps each of which can have a substantial effect on overall efficiency and performance. This section sets out the design rationale and chosen values for these processing steps, which was guided by the work of Navajas *et al.* [16].

#### 2.3.1. Signal conditioning & digitisation

This includes the amplification, filtering, sampling and analogue to digital conversion of the signal. An Intan RHD2132 was used to implement this step which provided the following options: a fixed gain of 192; tunable 3^*rd*^ order Butterworth low pass filter, single order analogue high pass filter and single order IIR digital high pass filter; up to 30 kSamples/s; and a 16-bit Analogue to Digital Converter (ADC). For action potential data the high pass corner was set to approximately 300 Hz (a balance between the modelled best setting for a high pass Butterworth filter, the band of interest and the need to reject power line interference), and a low pass corner of 3 kHz (to reduce high frequency noise and balance performance against the reduced data rate of sampling at a rate of 15 kSamples/s). Modelling had also shown that the sample depth could be reduced to as low as 7-bits without significantly impacting performance, in this work we converted the 16-bit signal to a 9-bit value to retain a higher dynamic range while reducing the data rate, processing requirements, and also to make efficient use of the memory structures of the FPGA used to perform the spike detection and template matching. The 9-bits to be used are user configurable at run time, but in practice the 9-bits chosen are rarely changed as the Most Significant Bits (MSBs) of the ADC are impractically large for neural signals and the noise floor of the front end is approximately 16 times the Least Significant Bit (LSB) of the ADC (2.4 *µ*V versus 0.195 *µ*V).

#### 2.3.2. Spike windowing

This step includes detection of the spikes, peak alignment and the windowing of the neural spike to be compared to the template. Spike detection is not essential for template matching – e.g. Schwarz *et al.* continuously test every incoming sample against templates – however, it can greatly reduce the amount of processing required in the template matching step and therefore increases system efficiency. This is a step that is highly parallel and was a key driver for utilising an FPGA (which can run many parallel calculations at a low clock speed) rather than a microcontroller (which would have required a high clock speed and hence high power consumption). Spike detection can be achieved by a variety of approaches, but simply detecting a threshold crossing remains an efficient and high performance solution [20] and was the chosen approach for this system with spike thresholds user configured on a per-channel basis (automatically calculated from the standard deviation of the median filtered noise [19]). The detected spikes were then peak aligned and a window of 5 samples before the peak and 10 after the peak (16 samples total) were passed to the template matching processing.

#### 2.3.3. Template Matching

This step includes the scoring of the differences between an incoming windowed spike snippet and a characteristic waveform (templates) of previously observed spikes, as well as the final classification of a spike. Navajas *et al.* reviewed numerous methods for comparing waveforms and there were 2 clear leading algorithms in the tradeoff between performance and efficiency – Norm 1 (sum of absolute difference) and Euclidean distance (sum of squared difference). In this instance (due to the lack of dedicated multiplication resource on the chosen FPGA) the Norm 1 implementation (shown in Figure 4) utilised Significantly less logic elements and was preferred.

**Figure 4.**
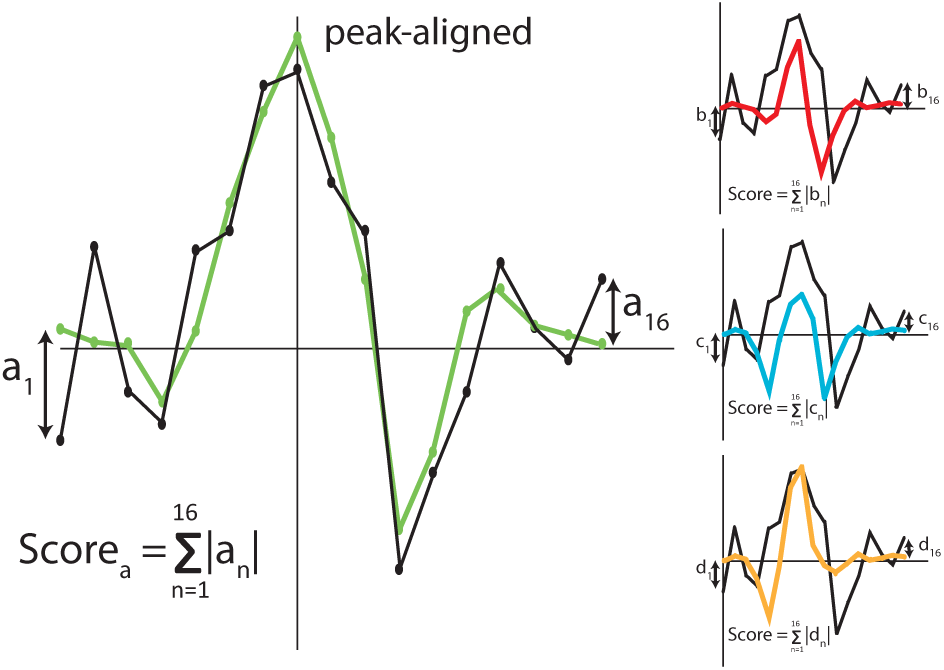
Calculation of template matching scores – an incoming spike waveform is compared against each of the templates and the sum of absolute difference for each pairing is calculated.

### 2.4. System Implementation

Figure 5 shows a block diagram breakdown of the key components across the 3 boards that make up the system. As shown the head-mounted boards can be powered either by a battery or from isolated USB power when tethered. When two power sources are available, a diode prevents uncontrolled current flowing into the battery and, due to its higher voltage, USB power is prioritised.

**Figure 5.**
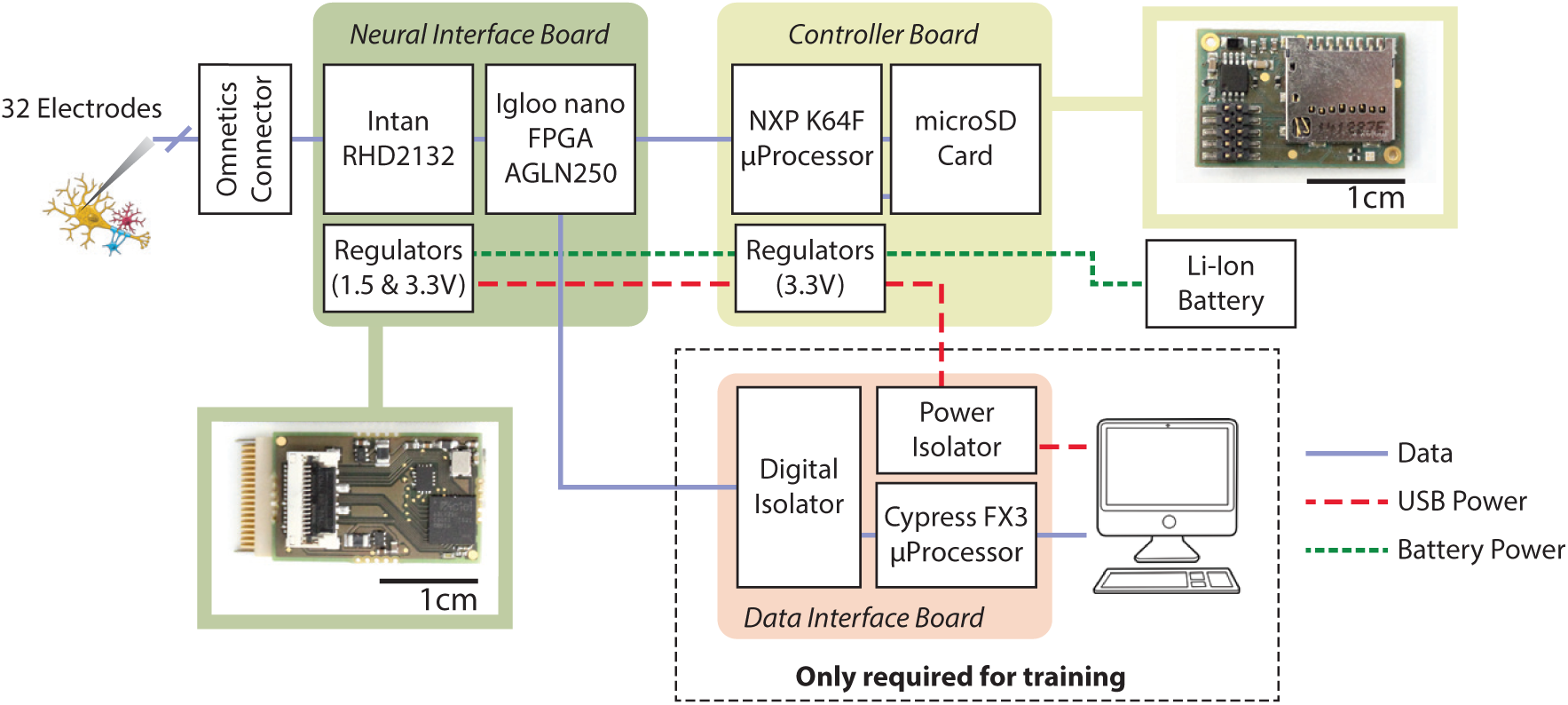
System architecture of the proposed platform with inset photos showing the corresponding head mounted modules.

#### 2.4.1. Data flow

The front end neural signal acquisition provides 32 channels of signal conditioning and digitisation. The data is streamed out to a small low power FPGA over an Serial Peripheral Interface (SPI) link. The FPGA performs signal manipulation to reduce the 16-bit sample to 9-bits before performing timestamping, and in normal operation it also performs spike detection and template matching (as depicted in Figure 6) before packaging the sorted result and/or raw neural signal data into 16-bit words for transmission. These words flow into one or both of the FPGA’s 2 output FIFO buffers – a large one feeding the wired SPI link (with the FPGA as slave), and a small one feeding the SPI link to the controller board (with the FPGA as master).

**Figure 6.**
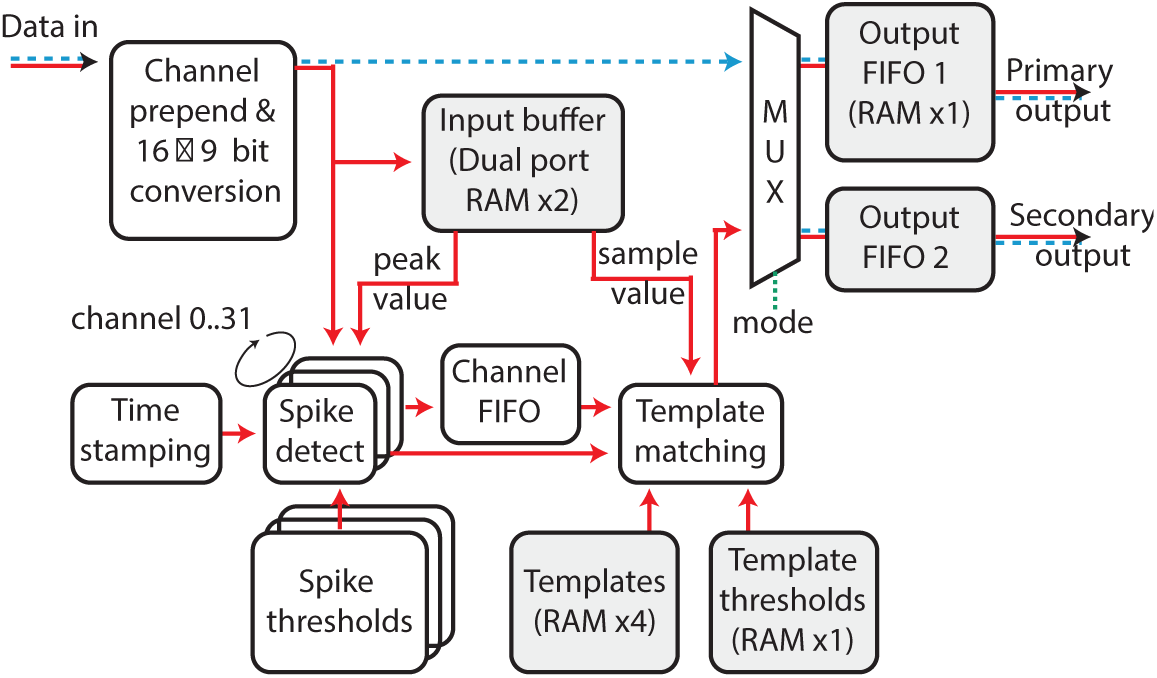
Data processing in the FPGA. The red line indicates data flow for template matching, while the blue dotted line indicates the data flow for raw neural signal data.

#### 2.4.2. Control & Communication

Control and configuration of the headstage is typically performed through the wired link during the calibration phase. While this upload is happening the microcontroller receives copies of these settings and backs them up to specifically named files on the SD card. Then if power is interrupted during normal untethered operation the microcontroller will read the configuration files and upload these values to the headstage restoring the programmed configuration. This also means that it is possible to operate the system completely without the wired link by simply putting appropriately coded files onto an SD card, inserting the card and power cycling the controller board. The full list of FPGA configurable registers are listed in Table 1, in addition to this are the settings required for the Intan front end (as specified on datasheet).

**Table 1.** configurable registers on FPGA

| Description | Bits | Scope | Total bits |
| --- | --- | --- | --- |
| Operation mode<sup>1</sup> | 4 | Global | 4 |
| LSB re-mapping<sup>2</sup> | 3 | Global | 3 |
| Sampling rate divider<sup>3</sup> | 4 | Global | 4 |
| Spike detection threshold | 9 | Channel | 288 |
| Templates | 576 | Channel | 18432 |
| Matching threshold | 64 | Channel | 2048 |

<sup>1</sup> Sets raw neural signal and/or sorted spike events output on the 2 links
<sup>2</sup> Controls the conversion of 16-bit samples to 9-bit
<sup>3</sup> Controls the global clock division.

#### 2.4.3. Clock control

Reducing clock speed is a key factor in improving system power performance. Here the sampling rate of the ADC and the two FPGA clock domains are derived by integer division (between 2-16) down from an off-chip clock running at 19.2 MHz. With nominal settings the sampling rate is at *∼*15 kS/s per channel (with an SPI clock speed of 9.6 MHz) and the FPGA domains are in turn divided down to 4.8 MHz and 2.4 MHz. All these frequencies are intrinsically linked (based on clock cycles necessary to process the data), so for efficient capture of low frequency Local Field Potentials (LFPs) the global integer division can be set to 16 giving a sampling speed of *∼*2 kS/s (SPI clock at 1.2 MHz) and clock domains of 600 kHz and 300 kHz.

#### 2.4.4. Data logging

In this instance the system has been equipped with a controller board for logging data. The core component of this board is a K64F microcontroller which implements three SPI interfaces (as shown in Figure 7) which receive data from the wired connection and from the FPGA and also enables control of the FPGA for recovery on power cycling.

**Figure 7.**
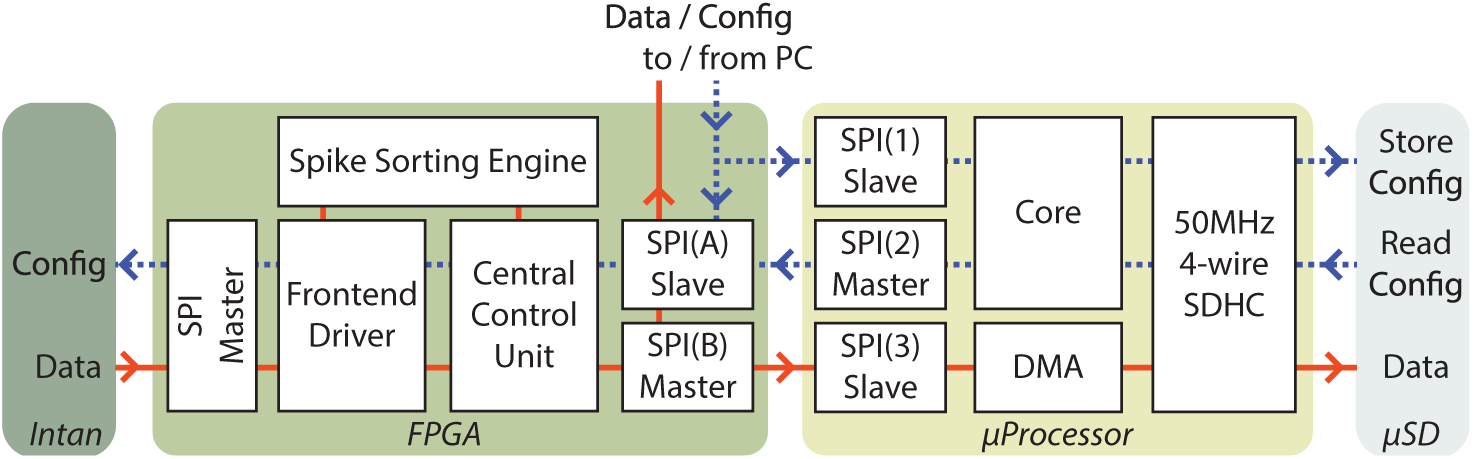
Data (red lines) and configuration (blue lines) flow between the head mounted modules.

The microcontroller receives data and uses a Direct Memory Access (DMA) channel to pass the data to the micro-SD card via an SDHC interface. For system validation purposes the aim was to record all raw and spike sorted data, as such the data output rate of the system was *∼*11 Mbps. This may not at first seem challenging (any class 10 SD card can write at 80 Mbps), however, the latency of typical SD cards can exceed 100 ms and presents a substantial challenge due to the data buffering requirements and limited memory on a microcontroller. As a result this system required careful code optimisation to maximise the memory available for buffering and align writing block size with sectors; and this combined with a U3 grade micro-SD card (with an average latency around 10 ms) ensured data integrity in repeated long term testing. Since the total amount of data generated for 24 hours is approximately 109 GB*‡*, a 128 GB card is required and the exFAT file system was used to enable large multi gigabyte files to be stored.

### 2.5. Computer interface

The system can be tethered to a PC using the Data Interface board which converts between an SPI interface and USB 3.0. A USB 3.0 interface is not required for 32 channels, however, the board was designed to be scalable by connecting up to 32 Neural Interface boards simultaneously in parallel – supporting up to 1,024 channels of raw and spike sorted data and increasing the total data rate to approximately 350 Mbps. In order to handle this high data rate, each bit sent over the 32 parallel MISO data lines from these boards is read in as a 4 byte word and this data is communicated directly to the USB interface using a DMA channel.

## 3. Results

### 3.1. System specification tests

The system was tested with benchtop equipments including a neural signal generator before *in-vivo* deployment. The hardware performance is summarised in Table 2.

**Table 2.**
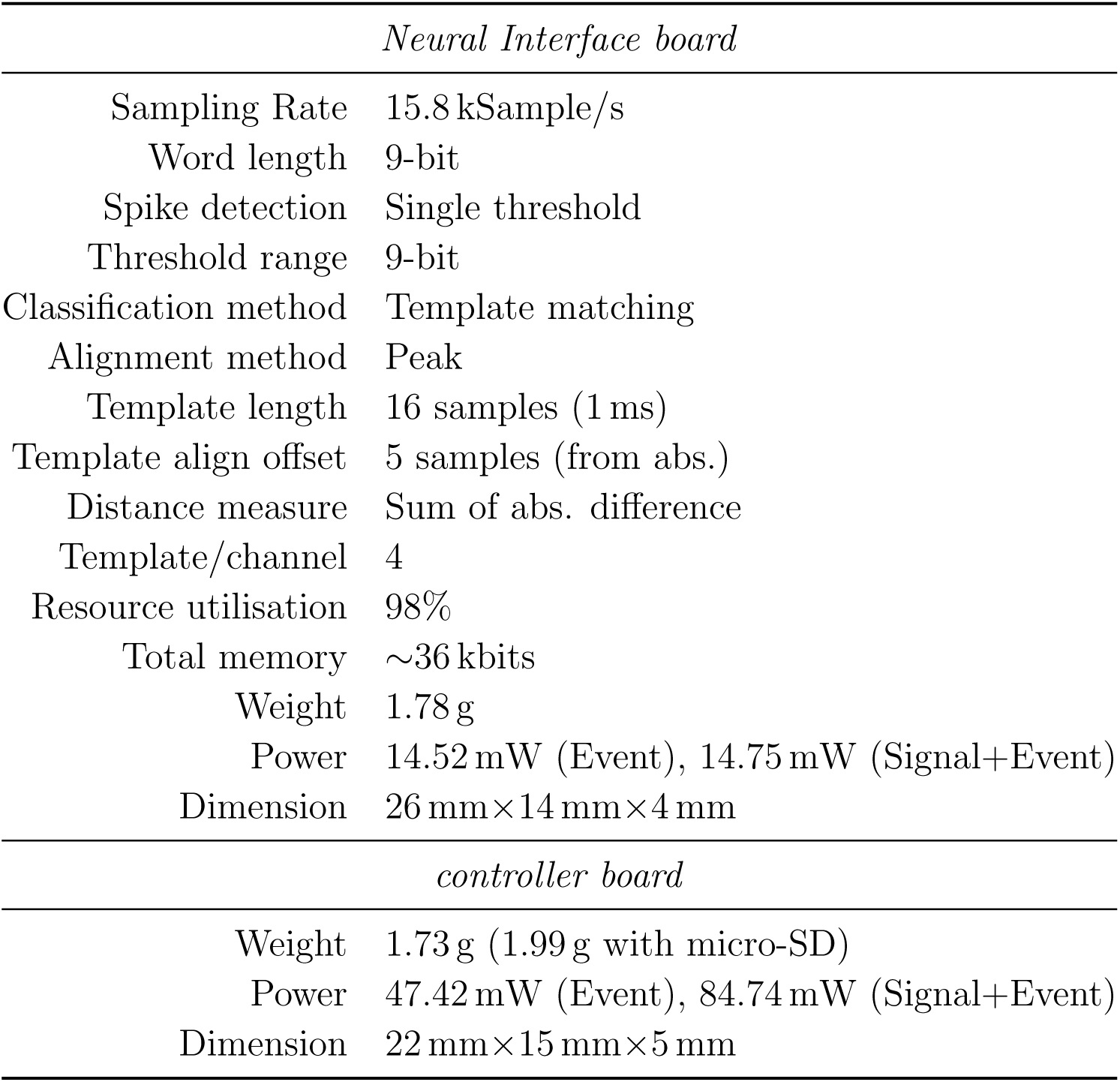
Specifications for key components

| *Neural Interface board* |  |
| --- | --- |
| Sampling Rate | 15.8 kSample/s |
| Word length | 9-bit |
| Spike detection | Single threshold |
| Threshold range | 9-bit |
| Classification method | Template matching |
| Alignment method | Peak |
| Template length | 16 samples (1 ms) |
| Template align offset | 5 samples (from abs.) |
| Distance measure | Sum of abs. difference |
| Template/channel | 4 |
| Resource utilisation | 98% |
| Total memory | ~36 kbits |
| Weight | 1.78 g |
| Power | 14.52 mW (Event), 14.75 mW (Signal+Event) |
| Dimension | 26 mm × 14 mm × 4 mm |
| *controller board* |  |
| Weight | 1.73 g (1.99 g with micro-SD) |
| Power | 47.42 mW (Event), 84.74 mW (Signal+Event) |
| Dimension | 22 mm × 15 mm × 5 mm |

‡ If logging all data including both the raw signal and sorted spike events.

Data integrity was tested by replacing the ADC’s output data with a simple counter on the FPGA – giving a known signal with the same data bandwidth as in real recording – and checking the sequence stored on the SD-card. A battery of 5200 mAh supports recording up to *∼*52 hours when recording both spike events and the raw neural signal. This differs from a simple calculation of battery capacity divided by measured power consumption due to the dropping battery output voltage over a discharge cycle.

### 3.2. In-vivo recording

The system was validated *in-vivo* in a rhesus macaque with moveable microwire electrodes (50 *µ*m tungsten, tip-impedence *∼*500 kΩ) [18] implanted into the motor cortex (M1) and protected with surgical cement and a skull mounted titanium case. A cradle to hold the platform was 3D-printed to fit inside the head case as shown in Figure 8. All experiments were approved by the local ethics committee and performed under appropriate UK Home Office licenses in accordance with the Animals (Scientific Procedures) Act 1986.

**Figure 8.**
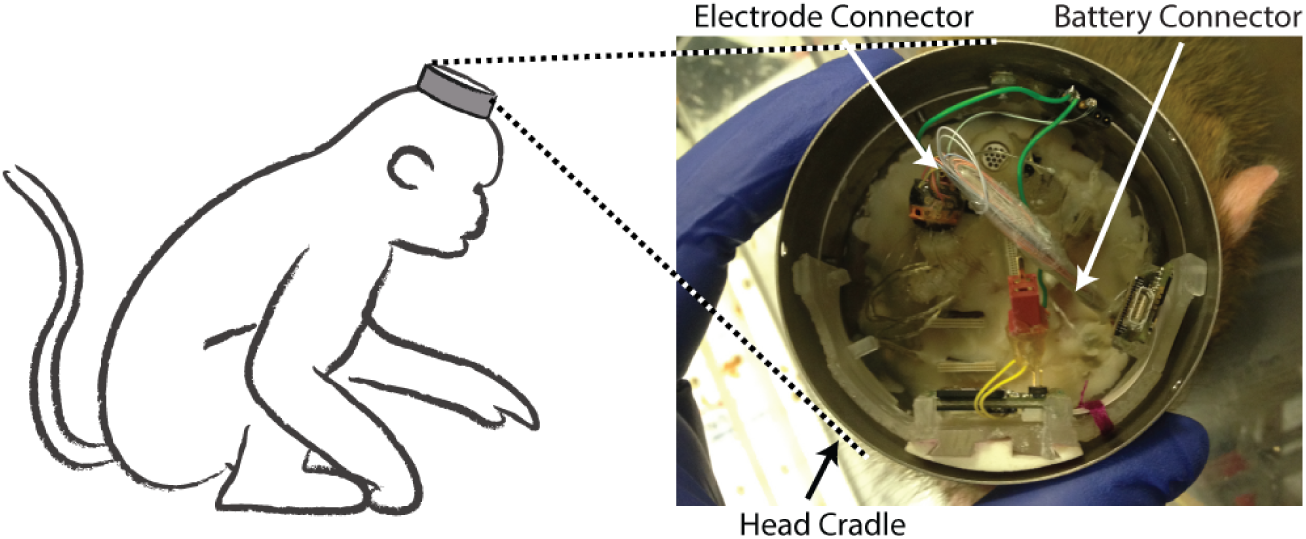
Experimental setup for the 24 hour recordings with a rhesus monkey. 12 tungsten wire electrodes are connected to the recording module. A cap with battery included was secured on top of the cradle.

Two 24-hour sessions were performed. In the first session, the system was configured to record only action potentials with on-line spike sorting. Once configured, the system was left recording for a 24 hour duration while the monkey was freely moving in her home cage. A total of 12 electrodes were connected across which 21 different cells were detected. The channel with the most units detected (#5) is shown in Figure 9. A two minute initial recording was used for clustering and 4 different templates were generated. Template 4 was merged into template 1 due to a close similarity. Figure 10 shows the resulting 3 spike templates (with the first 1,000 matched spikes) together with firing rates and inter-spike intervals (ISI) for channel #5 during the 24-hour session. The changes in firing rate during sleep can be clearly marked. It can however also be observed (from the ISI diagrams) that for this channel templates T1, T2 or T3 may contain more than one cell. An off-line analysis with Wave Clus yielded similar results demonstrating that this particular channel is challenging to sort for both online and offline sorters. Figure 11 shows a different recording channel (#3) where two units can be clearly identified.

**Figure 11.**
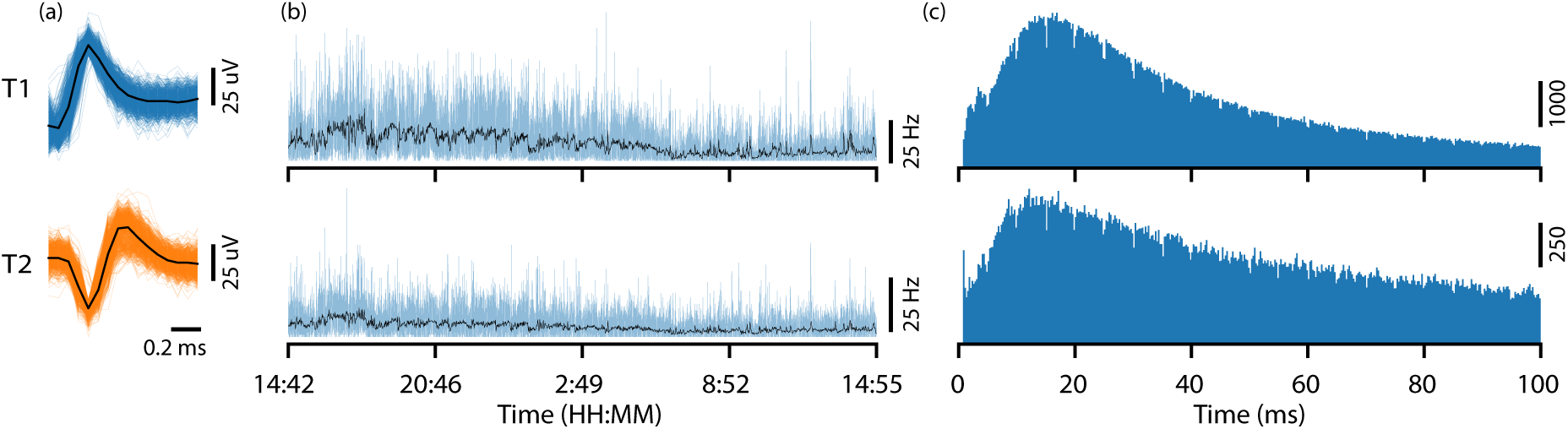
Example spike event data (for channel #3) generated the during the first 24 hour recording session. Shown are: (a) peak aligned and sorted first 1000 matched spikes and the characteristic templates; (b) spike firing rates averaged over 1 second (light blue) and 1 minute (black); and (c) Inter-Spike Interval (ISI).

**Figure 10.**
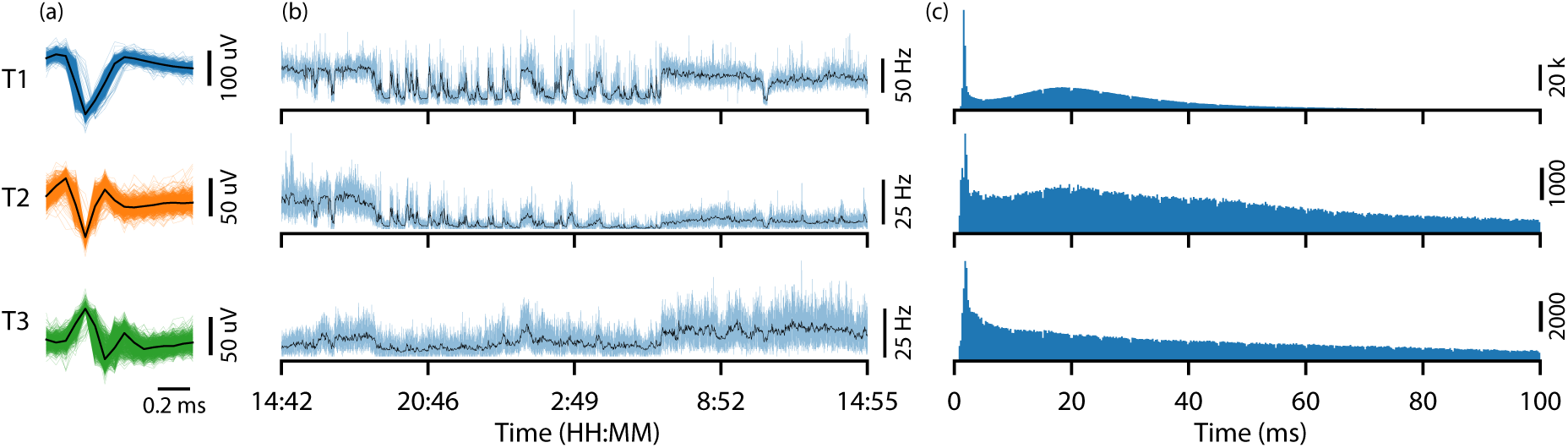
Example spike event data (for channel #5) generated the during the first 24 hour recording session. Shown are: (a) peak aligned and sorted first 1000 matched spikes and the characteristic templates; (b) spike firing rates averaged over 1 second (light blue) and 1 minute (black); and (c) Inter-Spike Interval (ISI).

**Figure 9.**
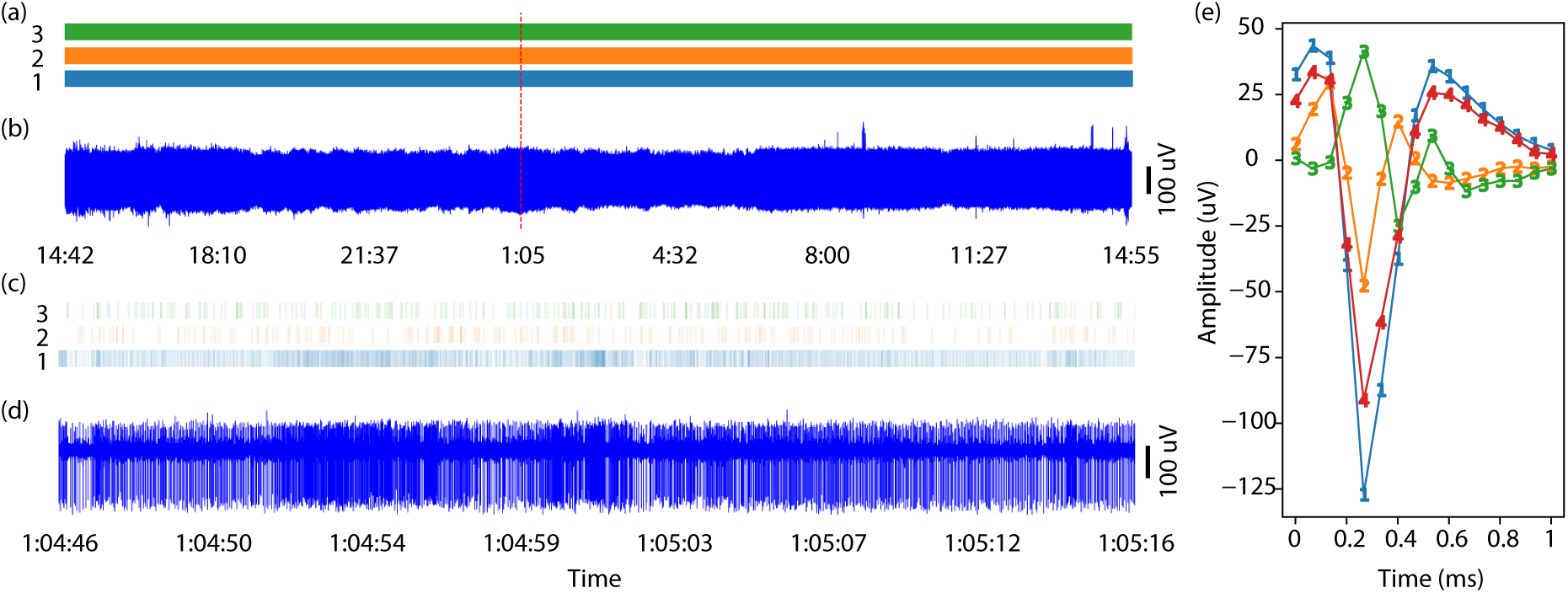
Extracellular neural recording (for channel #5) showing sorted spike events (in real-time) during the first 24 hour session. (a) and (b) are spike events and neural signal for the whole recordings showing consistent amplitude and constant operation throughout; (c) and (d) show a 30 s segment as marked by the dashed red line; Templates (e) generated using 2 min recording before the experiment using Wave Clus.

In the second session, the micro-SD card was replaced during a behavioural training session and the system was configured to record both action potentials and LFPs. The LFP signal and its spectrogram are shown in Figure 12. The increased power in the low frequency (*<*20 Hz) can be attributed to the activity expected during sleep. The hours of sleep are consistent with the period identified previously and individual sleep cycles can be observed.

**Figure 12.**
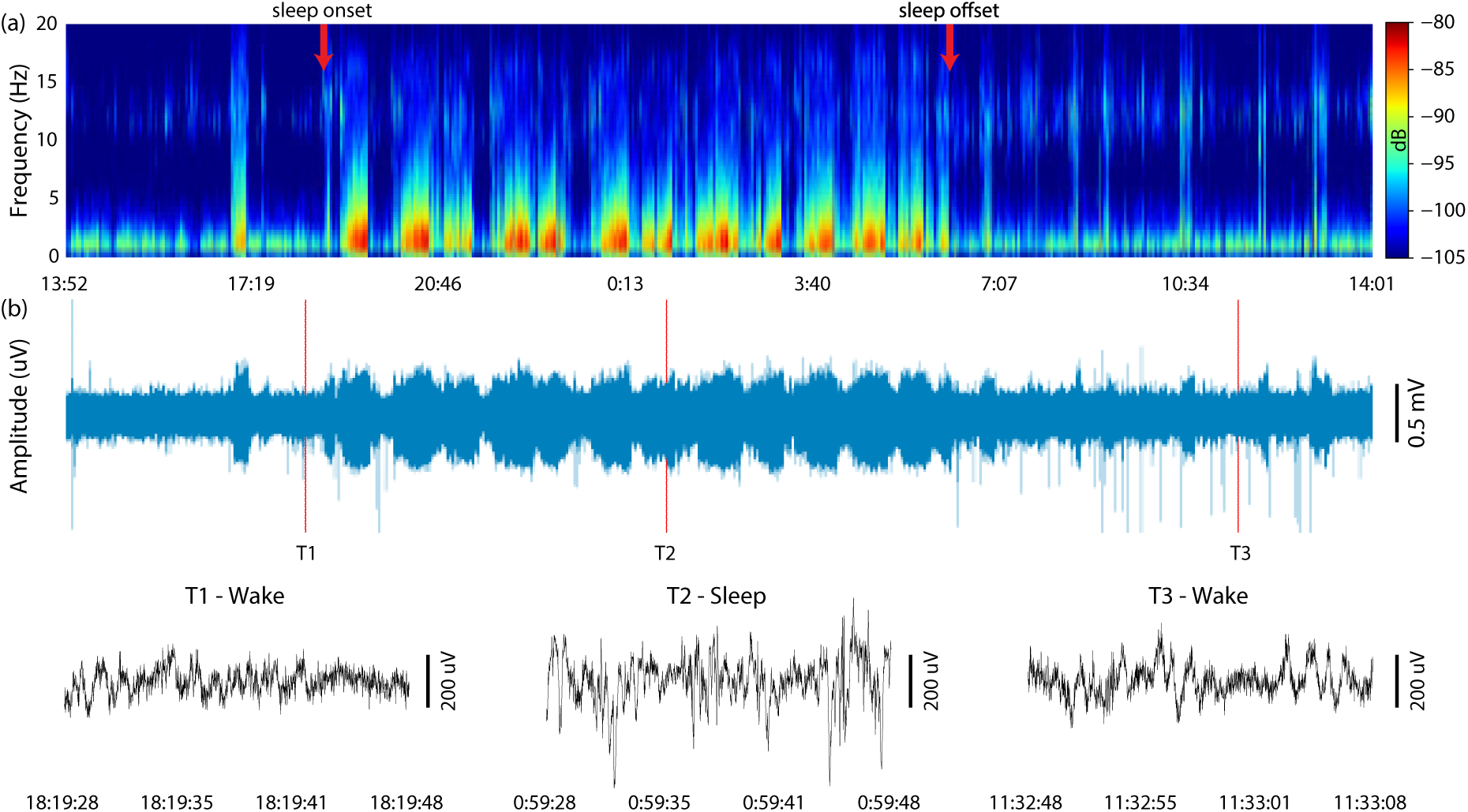
(a) Calculated spectrogram with sleep onset and cessation annotated; (b) Recorded LFP signal with magnified insets.

### 3.3. Power consumption

The power consumption was measured using an Agilent N6782A SMU (source measure unit) instrument. The idle power consumption was 52.83 mW (with SD card inserted) rising to a maximum of 99.45 mW when saving both the raw neural signal and spike event data to the SD card. The power when saving only spike events is 61.94 mW. A breakdown of the measured power budget is shown in Figure 13. Apart from the FPGA core which is running at 1.5 V, all other I/O and chips are running at 3.3 V. It should be noted that the idle current consumption is dominated by the microcontroller (*∼*82% of idle power) and that the power consumed by the spike sorting FPGA is minimal (*<*5.8% power in the worst case spike sorting only mode).

**Figure 13.**
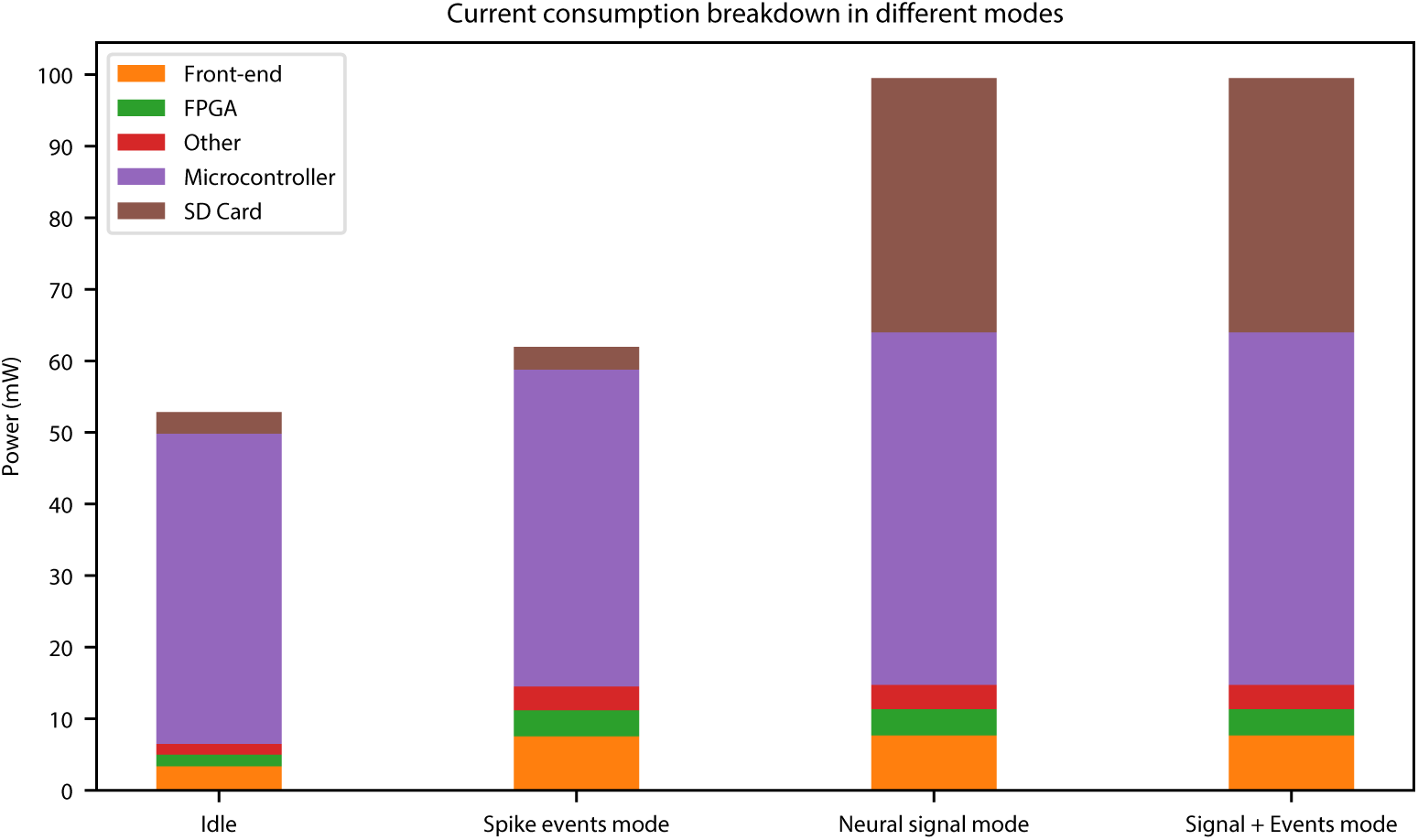
Breakdown of the current consumption for different modes of operation.

### 3.4. Data Reduction

The data storage required for recording the raw digitised neural signal for 24 hours for all 32-channels (excluding data encoding) is

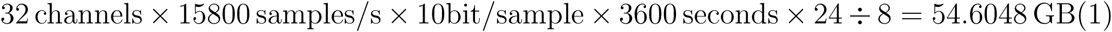

Whereas the actual recorded spike sorted data size was merely 66.3 MBytes giving a data reduction factor of approximately 823.

## 4. Discussion

The system presented here successfully demonstrated 24-hour on-node spike sorting and captured valuable data during untethered sleep showing characteristic patterns of reduced motor cortex spiking activity and patterns of modulation of LFP power spectral density This small and low power device enables massive data compression (and hence power reduction) while also providing triggers that could be invaluable for other event driven systems.

The spike sorting front end was paired in this instance with miniaturised SD-card logging due to a desire to reliably record neural signal and spike sorted data from our 32-channel headstage for validation purposes while minimising power consumption and hence battery size and weight for 24-hour trials. Our estimates of likely power consumption (see Figure 14) indicated that flash memory represented the lowest power off-the-shelf solution to our data logging needs and also demonstrated the Significant impact that spike sorting can have on system logging capability, power consumption and battery lifetime. Table 3 shows a comparison to similar platforms.

**Table 3.** Comparison with the State-of-the-Art

| Ref. | [units] | HermesD [21] | Neurochip-2 [22] | Schwarz et al. [1] | Willow [23] | This work |
| --- | --- | --- | --- | --- | --- | --- |
| # Channels | - | 32 | 3 | 512 | 1024 | 32 |
| Power/Ch. | [mW] | 4.43 | 109.33 | 2.06 | 6.84 | 1.94*, 3.11$\diamond$ |
| Data Storage | - | Wireless | SD ($\leq 2$ GB) | Wireless | SATA & Ethernet | SD |
| Spike sorting | - | No | TAWD | TM | No | TM |
| Auto-start | - | N/A | N/A | N/A | N/A | Yes |
| Size* | [mm] | 28×2×13 | 35×35×20 | 13.7×21.5×1.6†<br>43×21×3.4‡ | 150×160×20 | 26×14×4†<br>22×15×5‡ |
| Weight* | [g] | N/A | 145 | 11.6 | 164 | 1.99 |

\*Excl. battery, † Headstage, ‡ Interface, \*Spike Events, $\diamond$ Signal + Spike Events TM = Template Matching, TAWD = time-amplitude window discriminator

**Figure 14.**
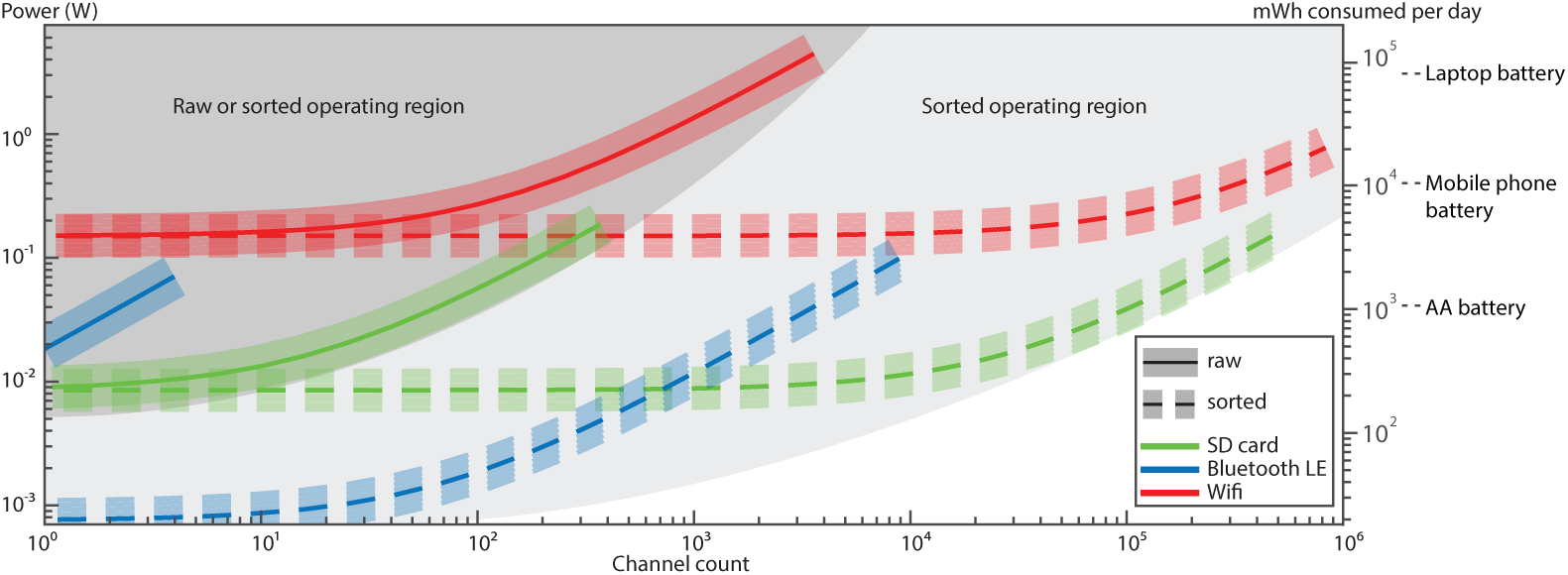
Estimate of power consumption for transmission (by Bluetooth Low Energy or WiFi 802.11g/ac) or local logging of data for exponentially increasing numbers of channels of neural signal or spike sorted data. Key assumptions include: a sampling rate of 16 kHz and an average spike rate of 10 spikes/s/channel. Wifi power estimates are based on a minimum range and low congestion scenario with *<*100 ms packet latency.

### 4.1. Expandability

Underpinning our design was the desire to create a device that rather than simply demonstrating technology, will actually be used on a day to day basis by widespread neuroscience groups. As such the system has been designed to be highly flexible and enables us to rapidly extend capability by adding, for example, a bidirectional wireless link or interfacing with a neuromodulation device. For instance, 2 Mbps supported by nRF52832 microcontroller is sufficient for streaming 7-channels of neural signal sampling at 16 kS/s with a resolution of 10-bit or 79-channels of spike events firing at 10 Hz. The microcontroller can also further processed the spike events or raw neural signal.

### 4.2. Recording duration

For compact system like the one presented here, its maximum continuous operating hours is typically limited by the capacity of the battery and/or the data storage media. This is a complex function of recording mode, number of channels enabled, filter and ADC settings and spike rate. In our case, for all 32-channels used for continuous recording in signal + events mode, a maximum 50.97 hours was measured. The total data storage required is 185.5 GB which doesn’t require a change of SD card. If all 32 channels operate records only spike events, the benchmarks are projected to 81.87 hours continuous recording and 225 MB of data based on the power and data reduction ratio reported in Section 3.

In both cases, the 5200 mAh battery used was not fully discharged but the system stopped at 3.65 V due to the required overhead of our regulators. If high efficiency DC-DC regulators are used and a 400 GB micro-SD card (the highest capacity currently available), then maximum continuous recording time for 32-channels in signal + events modes will be *∼*107 hours (limited by data storage) and *∼*156.8 hours (limited by battery) respectively. If the system is however operated in spike events mode (i.e. recording, spike sorting, and storing only the classified spike event times), both the system power consumption (Fig. 13) and memory requirements are Significantly reduced, resulting in a continuous operating time of *∼*252 hours (around 10.5 days) that is limited only by battery.

### 4.3. Future work

Technological advances mean that there are already substantial improvements we will be making to our system design. In particular it is worth noting that all the spike detection and sorting in this system was performed on a small (5 mm *×* 5 mm), low power and relatively old technology (130 nm) FPGA, however, technology scaling offers Significant potential to increase the channel count while also reducing the power consumption and footprint. Indeed we have since successfully ported the code to an FPGA family offering 6 times smaller footprint, 30 times as much memory (a key limiting factor on channel count) and 3 times lower core power consumption. Moving to an Application Specific Integrated Circuit (ASIC) design could offer a further step change in system performance and capability. Similarly the microcontroller core power consumption tracks with the technology and reducing the current consumption to a small fraction of its current level is feasible.

Changes to the front end (signal conditioning) ASIC are another aspect with potential for system-wide impact [24, 25]. The current front end is designed for generic biopotential recording and as a result its core power consumption is unnecessarily high (for this application). Power consumption could also be greatly reduced by moving the spike detection into the front end as this could reduce the data rate output at this stage by around 2 orders of magnitude [26, 27]. Performing spike detection was a key motivator for choosing an FPGA (due to the parallel nature of spike detection) and utilised around 50% of the FPGA resource, therefore moving spike detection to the front end potentially enables efficient microcontroller based spike sorting or reduces the FPGA requirements and required clock speed.

In the near term we are looking at a variety of algorithm enhancements that could improve template matching performance (e.g. by pre-manipulation of the templates [28]) or identify and potentially track changes in templates due to electrode drift and fibrous encapsulation – reducing calibration requirements and enhancing system performance for chronic recordings. While in the long term we would look to leverage commercial technology scaling to enable integration of on-line naive spike sorting for hundreds of channels – scaling, enhancing and making viable existing approaches [29, 30].

## 5. Conclusion

Challenges abound for delivering the next generation of neural interface technology. As the number of recording channels increases by orders of magnitude so too will the data rate as well as the power required for data transmission and the storage requirements. It is therefore clear that simply scaling existing electrode arrays, connectors, electronic designs or wireless bandwidth will not deliver the performance required in future and that fundamentally new approaches are required. Spike sorting on the node is a key enabling capability in dealing with the coming data barrage and for creating untethered BMIs and closed loop event driven neuromodulation. The system presented here represents our first efforts to deploy this capability into the neuroscience community.

## Acknowledgement

This research is funded by Engineering and Physical Sciences Research Council (EPSRC) EP/I000569/1, EP/H051570/1, EP/H051651/1, EP/K015060/1. We thank Dr. Jennifer Tulip and Dr. Wei Xu from Newcastle University for their support on *in-vivo* experiments and data analysis.

## References

[1] Schwarz D A, Lebedev M A, Hanson T L, Dimitrov D F, Lehew G, Meloy J, Rajangam S, Subramanian V, Ifft P J, Li Z et al. 2014 Chronic, wireless recordings of large-scale brain activity in freely moving rhesus monkeys Nature Methods 11 670–676

[2] Yeon P, Mirbozorgi S, Ash B, Eckhardt H and Ghovanloo M 2016 Fabrication and Microassembly of a mm-Sized Floating Probe for a Distributed Wireless Neural Interface Micromachines 7 154 ISSN 2072-666X

[3] Ando H, Takizawa K, Yoshida T, Matsushita K, HirataMand Suzuki T 2016 Wireless Multichannel Neural Recording With a 128-Mbps UWB Transmitter for an Implantable Brain-Machine Interfaces IEEE Transactions on Biomedical Circuits and Systems 10 1068–1078 ISSN 1932-4545

[4] Bargmann C, Newsome W, Anderson A, Brown E, Deisseroth K, Donoghue J, MacLeish P, Marder E, Normann R, Sanes J et al. 2014 Brain 2025: a scientific vision Brain Research Through Advancing Innovative Neurotechnologies (BRAIN) Working Group Report to the Advisory Committee to the Director, NIH. Available online at: http://www.nih.gov/science/brain/2025/(US National Institutes of Health, 2014)

[5] Liu X, Zhang M, Xiong T, Richardson A G, Lucas T H, Chin P S, Etienne-Cummings R, Tran T D, Van Der Spiegel J, Liu X, Van Der J, Richardson A G and Lucas T H 2016 A Fully Integrated Wireless Compressed Sensing Neural Signal Acquisition System for Chronic Recording and Brain Machine Interface IEEE Transactions on Biomedical Circuits and Systems 10

[6] Aziz J N, Abdelhalim K, Shulyzki R, Genov R, Bardakjian B L, Derchansky M, Serletis D and Carlen P L 2009 256-channel neural recording and delta compression microsystem with 3D electrodes IEEE Journal of Solid-State Circuits 44 995–1005

[7] Kamboh A M, Raetz M, Oweiss K G and Mason A 2007 Area-power efficient VLSI implementation of multichannel DWT for data compression in implantable neuroprosthetics IEEE Transactions on Biomedical Circuits and Systems 1 128–135

[8] Chae M S, Yang Z, Yuce M R, Hoang L and Liu W 2009 A 128-channel 6mW wireless neural recording IC with spike feature extraction and UWB transmitter IEEE Transactions on Neuralystems and Rehabilitation Engineering 17 312–321

[9] Karkare V, Gibson S and Markovic D 2011 A 130μW, 64-channel neural spike-sorting DSP chip IEEE Journal of Solid-State Circuits 46 1214–1222

[10] Rizk M, Obeid I, Callender S H and Wolf P D 2007 A single-chip signal processing and telemetry engine for an implantable 96-channel neural data acquisition system Journal of Neural Engineering 4 309

[11] Eftekhar A, Paraskevopoulou S E and Constandinou T G 2010 Towards a next generation neural interface: Optimizing power, bandwidth and data quality Biomedical Circuits and Systems Conference (BioCAS), 2010 IEEE (IEEE) pp 122–125

[12] Quian Quiroga R 2012 Spike sorting Current Biology 22 R45–R46

[13] Rey H G, Pedreira C and Quian Quiroga R 2015 Past, present and future of spike sorting techniques Brain Research Bulletin 119 106–117

[14] Kamboh A M and Mason A J 2013 Computationally efficient neural feature extraction for spike sorting in implantable high-density recording systems IEEE Transactions on Neural Systems and Rehabilitation Engineering 21 1–9

[15] Barsakcioglu D Y and Constandinou T G 2016 A 32-channel mcu-based feature extraction and classification for scalable on-node spike sorting Circuits and Systems (ISCAS), 2016 IEEE International Symposium on (IEEE) pp 1310–1313

[16] Navajas J, Barsakcioglu D Y, Eftekhar A, Jackson A, Constandinou T G and Quian Quiroga R 2014 Minimum requirements for accurate and efficient real-time on-chip spike sorting Journal of Neuroscience Methods 230 51–64

[17] Williams I, Luan S, Jackson A and Constandinou T G 2015 A scalable 32-channel neural recording and real-time FPGA based spike sorting system Biomedical Circuits and Systems Conference (BioCAS), 2016 IEEE pp 188–191

[18] Jackson A, Mavoori J and Fetz E E 2007 Correlations between the same motor cortex cells and arm muscles during a trained task, free behavior, and natural sleep in the macaque monkey Journal of neurophysiology 97 360–374

[19] Quian Quiroga R, Nadasdy Z and Ben-Shaul Y 2004 Unsupervised spike detection and sorting with wavelets and superparamagnetic clustering Neural Computation 16 1661–1687

[20] Obeid I and Wolf P D 2004 Evaluation of spike-detection algorithms fora brain-machine interface application IEEE Transactions on Biomedical Engineering 51 905–911

[21] Miranda H, Gilja V, Chestek C a, Shenoy K V and Meng T H 2010 HermesD: a high-rate long-range wireless transmission system for simultaneous multichannel neural recording applications IEEE Transactions on Biomedical Circuits and Systems 4 181–191 ISSN 1932-4545

[22] Zanos S, Richardson A G, Shupe L, Miles F P and Fetz E E 2011 The Neurochip-2: an autonomous head-_xed computer for recording and stimulating in freely behaving monkeys. IEEE Transactions on Neural Systems and Rehabilitation Engineering 19 427–35 ISSN 1558-0210

[23] Kinney J P, Bernstein J G, Meyer A J, Barber J B, Bolivar M, Newbold B, Scholvin J, Moore-Kochlacs C, Wentz C T, Kopell N J and Boyden E S 2015 A direct-to-drive neural data acquisition system Frontiers in Neural Circuits 9 ISSN 1662-5110

[24] Barsakcioglu D Y, Eftekhar A and Constandinou T G 2013 Design optimisation of front-end neural interfaces for spike sorting systems Circuits and Systems (ISCAS), 2013 IEEE International Symposium on (IEEE) pp 2501–2504

[25] Barsakcioglu D Y, Liu Y, Bhunjun P, Navajas J, Eftekhar A, Jackson A, Quian Quiroga R and Constandinou T G 2014 An analogue front-end model for developing neural spike sorting systems IEEE Transactions on Biomedical Circuits and Systems 8 216–227

[26] Luan S, Liu Y, Williams I and Constandinou T G 2016 An event-driven soc for neural recording Biomedical Circuits and Systems Conference (BioCAS), 2016 IEEE (IEEE) pp 404–407

[27] Liu Y, Luan S, Williams I, Rapeaux A and Constandinou T G 2017 (accepted, in press) A 64-channel versatile neural recording SoC with activity dependent data throughput IEEE Transactions on Biomedical Circuits and Systems

[28] Frehlick Z, Williams I and Constandinou T G 2016 Improving neural spike sorting performance using template enhancement Biomedical Circuits and Systems Conference (BioCAS), 2016 IEEE (IEEE) pp 524–527

[29] Rutishauser U, Schuman E M and Mamelak A N 2006 Online detection and sorting of extracellularly recorded action potentials in human medial temporal lobe recordings, in vivo Journal of neuroscience methods 154 204–224

[30] Karkare V, Gibson S and Markovic D 2013 A 75_W, 16-Channel Neural Spike-Sorting Processor With Unsupervised Clustering IEEE Journal of Solid-State Circuits 48 2230–2238 ISSN 0018-9200

